## Supplementary Information for "Temperature Increase Drives Critical Slowing Down of Fish Ecosystems"

### 1 Supplementary Results

As proven in previous paper (Li and Convertino, 2021) we provide a broader analysis of species interaction inference results for different TE-estimator models. Fig. S5 shows how the Gaussian estimator provides a quite different pattern of interactions than the other three models (TE with Kernel estimator, CCM and the Pearson correlation coefficient). This is likely because the Gaussian estimator is based on a linearity assumption, meaning that it assumes linear interactions between variables (Lizier, 2014). The pearson correlation coefficient is a symmetrical measure of linear correlation between two sets of data. This measure only captures linear relationship between variables, it is not capable of distinguishing directed interactions, either. Even though $\rho$ from CCM and TE from OIF with Kernel estimator present similar patterns, the OIF-inferred heat map from Kernel presents larger gradient of interactions that highlight the divergence in fish populations of species 4-9 from other species compared to the heat map from CCM. This divergence in the distribution of species populations can be observed in Fig. S2. ~~It is worth noting that species 4-9 are all native species (see Table S1).~~ Therefore, OIF with Kernel estimator allows a better identification of species clusters considering gradients of inferred interactions. Additionally, TE from Kernel estimator is able to estimate some weak observed interactions such as of species 2 with others, while CCM essentially consider null interactions for these species. Therefore, the Kernel estimator was selected as the TE estimator in the OIF model (Li and Convertino, 2021).

### References

Jie Li and Matteo Convertino. Inferring ecosystem networks as information flows. Scientific Reports, 11(1):1–22, 2021.

Joseph T. Lizier. Jidt: An information-theoretic toolkit for studying the dynamics of complex systems. Frontiers in Robotics and AI, 1:11, 2014. ISSN 2296-9144. doi: 10.3389/frobt.2014.00011. URL https://www.frontiersin.org/article/10.3389/frobt.2014.00011.

| **Species** | **Fish Stock** | **Native/Invasive** | **Conservation Status** | **Other Use** | **Threat** |
|---|---|---|---|---|---|
| 1. *Aurelia aurita* (Moon jellyfish) | Yes | Native | NE | Ornamental | Venomous |
| 2. *Engraulis japonicus* (Japanese anchovy) | Yes | Native | LC | Aquaculture/Game | - |
| 3. *Plotosus lineatus japonicus* (Sea catfish) | No | Invasive | NE | - | Venomous |
| 4. *Sebastes inermis* (Black snapper) | Yes | Native | LC | Game | - |
| 5. *Trachurus japonicus* (Horse mackerel) | Yes | Native | NT | Aquaculture | - |
| 6. *Girella punctata* (Blackeye seabream) | Yes | Native | NE | Game | - |
| 7. *Pseudolabrus sieboldi* (Wrasse) | Yes | Native | LC | - | - |
| 8. *Halichoeres poecilopterus* (Rainbow wrasse) | Yes | Native | LC | Aquaculture/Game/Ornamental | - |
| 9. *Halichoeres tenuispinnis* (Chinese wrasse) | No | Invasive | LC | Ornamental | - |
| 10. *Chaenogobius gulosus* (Goby) | No | Native | NE | - | - |
| 11. *Pterogobius zonoleucus* (Blue/Yellow striped Goby) | No | Native | LC | - | - |
| 12. *Tridentiger trigonocephalus* (Chameleon Goby) | No | Native | NE | - | - |
| 13. *Siganus fuscescens* (Rabbitfish) | Yes | Invasive | LC | Aquaculture | Venomous |
| 14. *Sphyraena pinguis* (Red barracuda) | Yes | Native | NE | - | - |
| 15. *Rudarius ercodes* (Pigmy filefish) | No | Invasive | LC | - | - |

Table S1:

| **Species** | **OTE** | **ITE** | **Mean** | **Std** |
|---|---|---|---|---|
| 1. *Aurelia aurita* | 6.477 | 9.612 | 23.826 | 126.050 |
| 2. *Engraulis japonicus* | 8.158 | 3.412 | 68.056 | 379.300 |
| 3. *Plotosus lineatus japonicus* | 7.058 | 3.174 | 28.253 | 133.030 |
| 4. *Sebastes inermis* | 6.160 | 10.524 | 31.874 | 50.512 |
| 5. *Trachurus japonicus* | 6.393 | 8.029 | 174.940 | 258.700 |
| 6. *Girella punctata* | 6.051 | 7.767 | 14.863 | 29.911 |
| 7. *Pseudolabrus sieboldi* | 6.273 | 10.308 | 7.333 | 6.970 |
| 8. *Halichoeres poecilopterus* | 6.128 | 6.978 | 7.565 | 11.755 |
| 9. *Halichoeres tenuispinnis* | 6.660 | 5.789 | 17.575 | 33.523 |
| 10. *Chaenogobius gulosus* | 8.048 | 2.539 | 8.939 | 51.713 |
| 11. *Pterogobius zonoleucus* | 6.230 | 6.839 | 18.542 | 77.554 |
| 12. *Tridentiger trigonocephalus* | 6.986 | 11.227 | 31.458 | 43.289 |
| 13. *Siganus fuscescens* | 7.071 | 3.768 | 4.646 | 25.866 |
| 14. *Sphyraena pinguis* | 8.165 | 2.369 | 8.761 | 50.779 |
| 15. *Rudarius ercodes* | 6.005 | 9.527 | 12.142 | 33.463 |

Table S2:

| **Salience** | **All Temp.** | **≤10°C** | **10-15°C** | **15-20°C** | **20-25°C** | **≥25°C** |
|---|---|---|---|---|---|---|
| 1 | 7→10 (0.87) | 15→3 (0.33) | 15→3,12→13 (0.87) | 7→13 (0.73) | 7→14 (0.93) | 6→1,7→13 (0.87) |
| 2 | 5→2 (0.73) | 15→6 (0.27) | 7→5,7→8,11→9 (0.80) | 5→10 (0.67) | 7→3,12→10 (0.87) | 15→2 (0.80) |
| 3 | 5→1,7→13,7→14 (0.67) | 12→1,12→2,15→4 (0.20) | 7→1,7→4,12→6,11→10 (0.73) | 7→3 (0.53) | 7→2,7→15 (0.80) | 7→3 (0.6) |
| 4 | 7→3,7→4,7→9,7→11 (0.53) | Many (0.13) | 7→15 (0.67) | 7→11 (0.33) | 7→11 (0.67) | 5→12 (0.53) |
| 5 | 7→15 (0.40) | Many (0.07) | 12→2,12→11 (0.40) | 6→1,5→2,5→7 (0.27) | 7→6 (0.53) | 7→10 (0.47) |

Table S3:

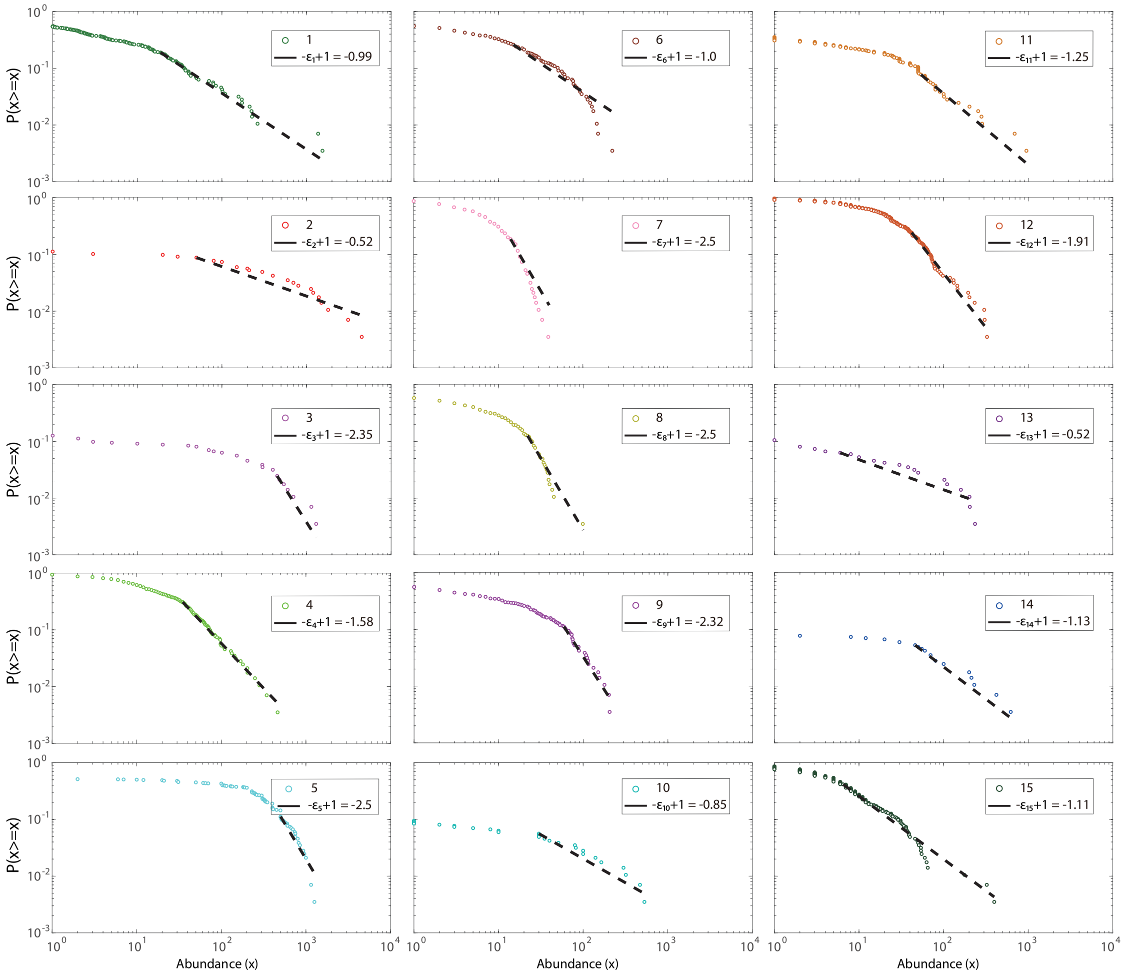
P(x>=x)
Abundance (x)
1
$-\epsilon_1+1 = -0.99$
2
$-\epsilon_2+1 = -0.52$
3
$-\epsilon_3+1 = -2.35$
4
$-\epsilon_4+1 = -1.58$
5
$-\epsilon_5+1 = -2.5$
6
$-\epsilon_6+1 = -1.0$
7
$-\epsilon_7+1 = -2.5$
8
$-\epsilon_8+1 = -2.5$
9
$-\epsilon_9+1 = -2.32$
10
$-\epsilon_{10}+1 = -0.85$
11
$-\epsilon_{11}+1 = -1.25$
12
$-\epsilon_{12}+1 = -1.91$
13
$-\epsilon_{13}+1 = -0.52$
14
$-\epsilon_{14}+1 = -1.13$
15
$-\epsilon_{15}+1 = -1.11$

Figure S1:

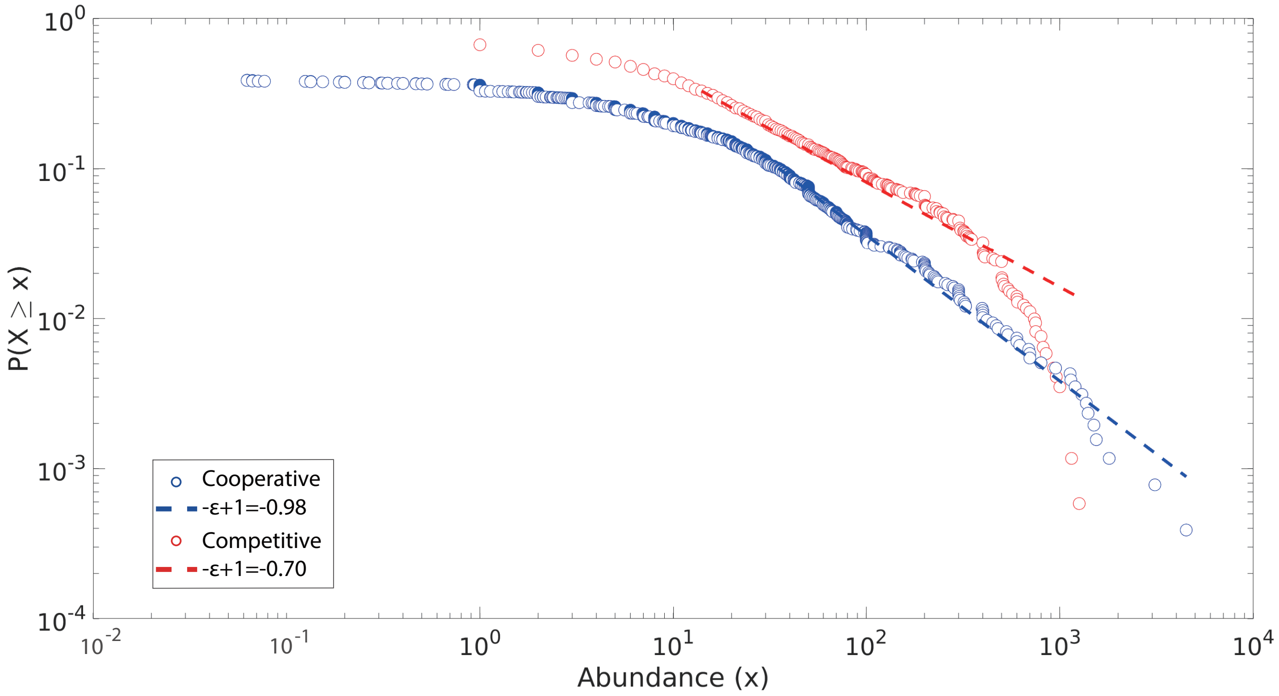

Figure S2:

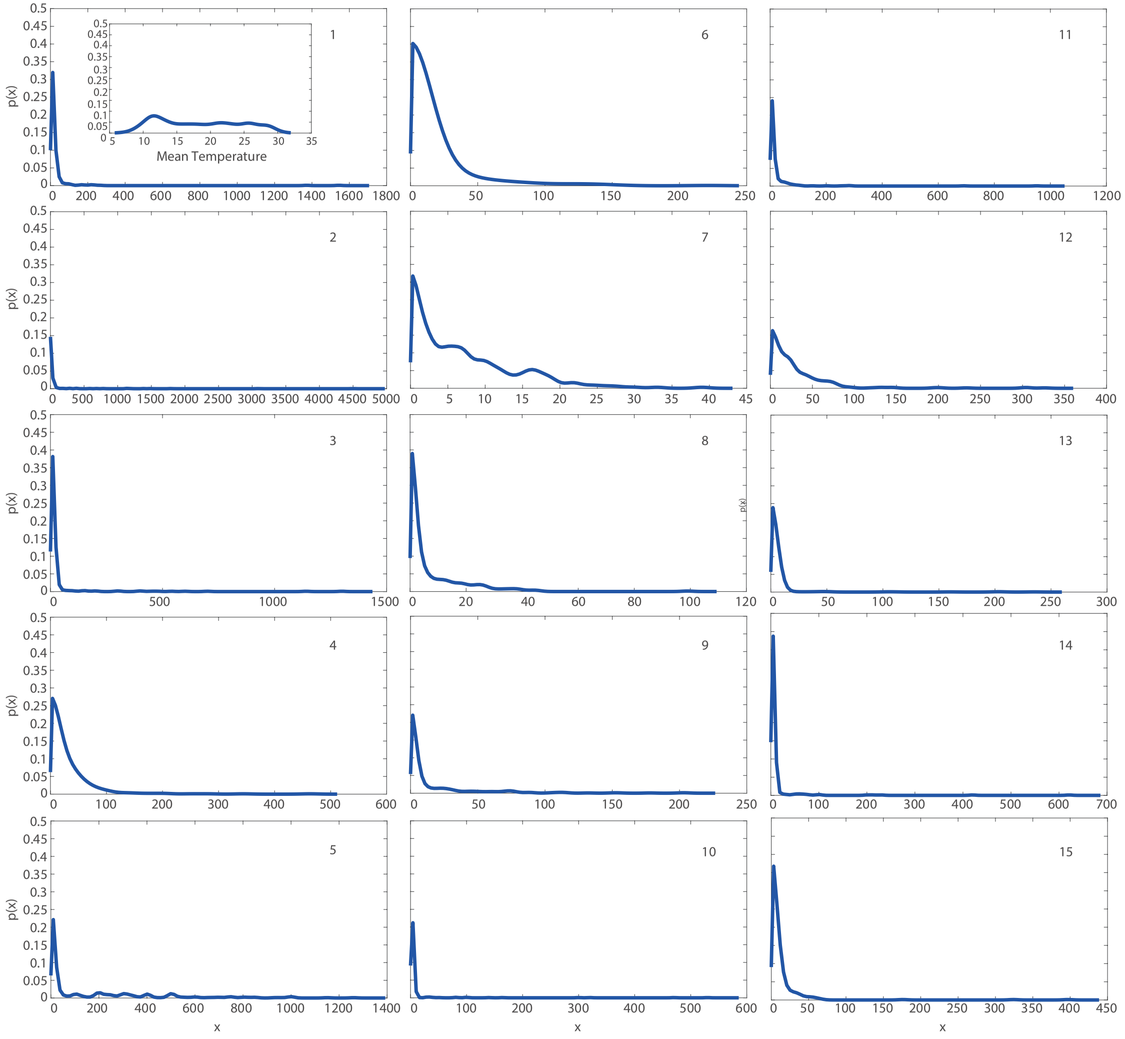

Figure S3:

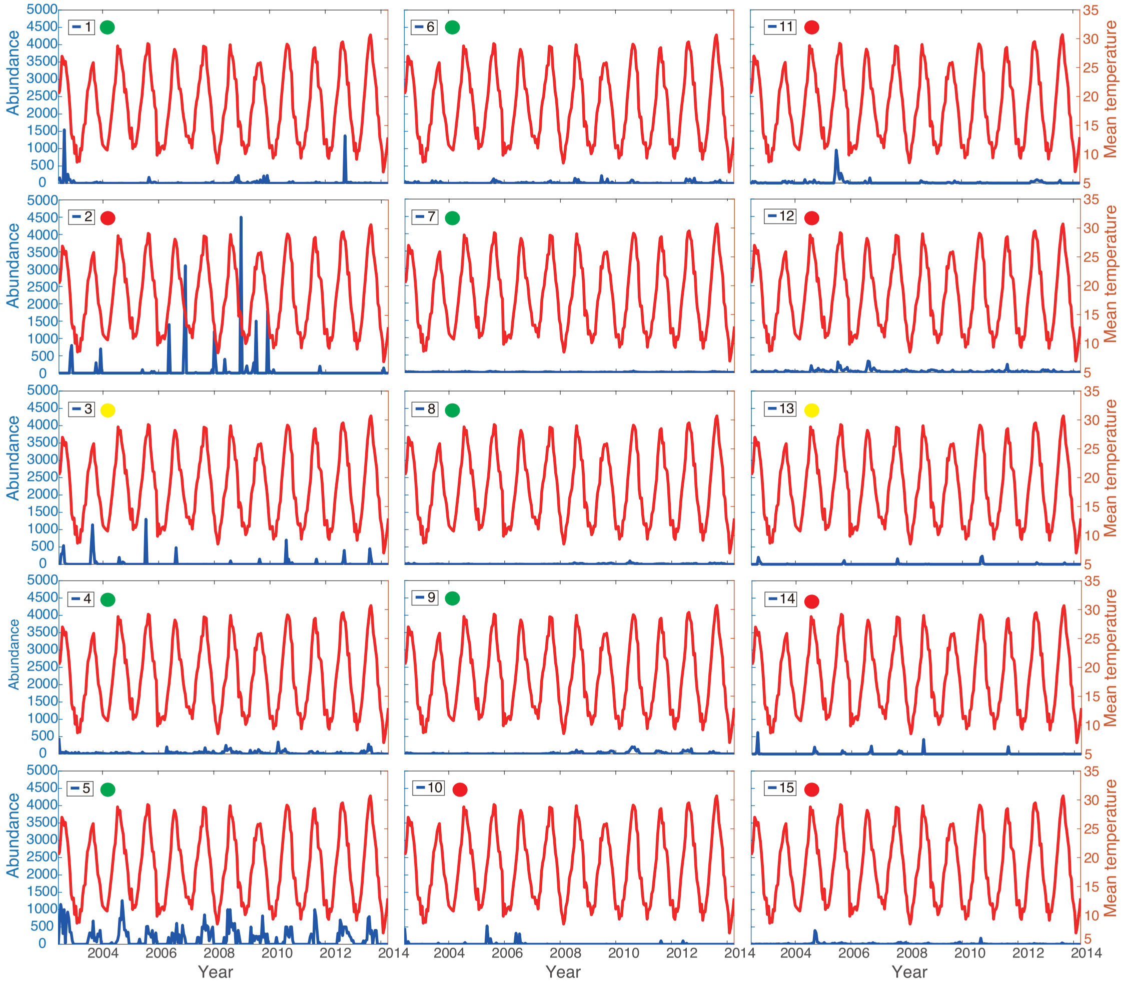

Figure S4:

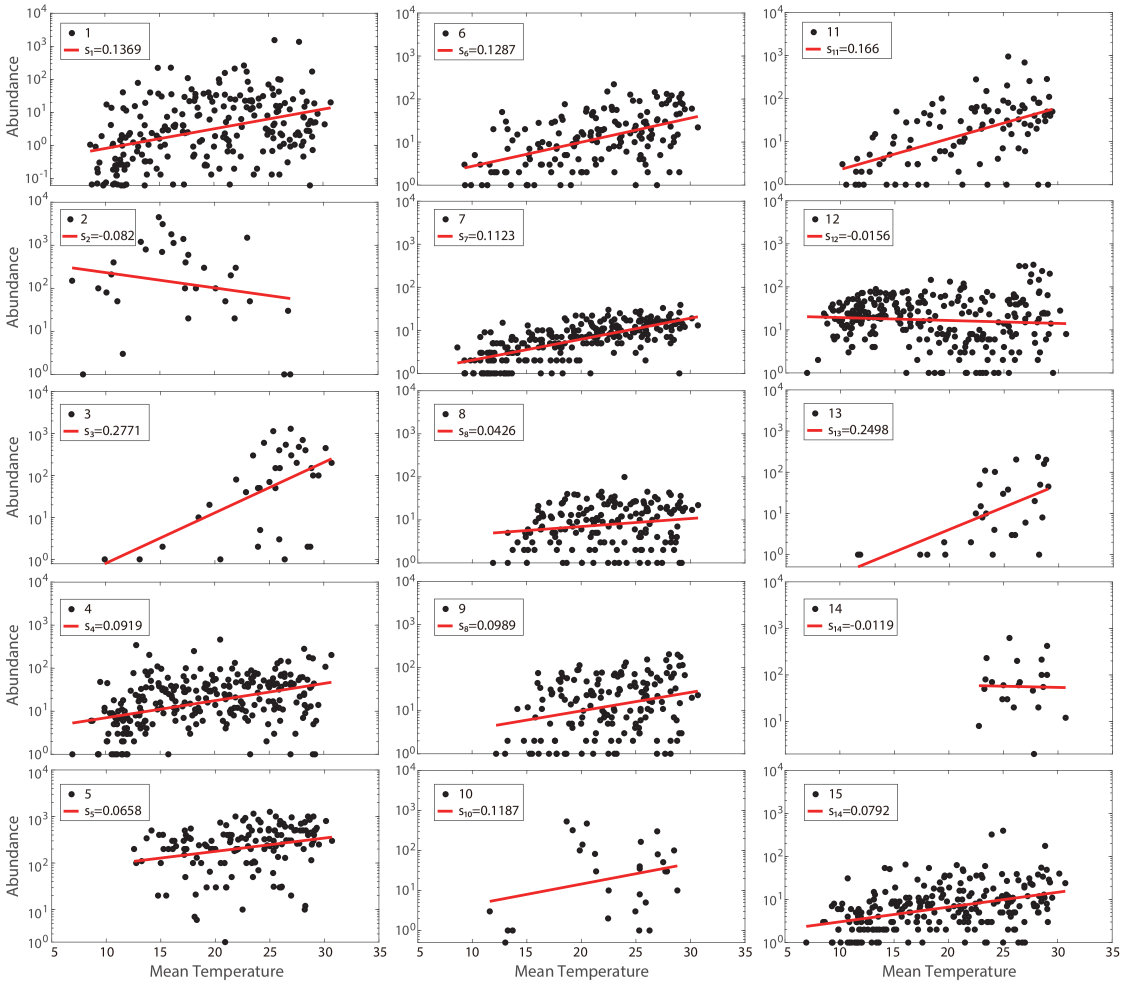

1
s1=0.1369
2
s2=-0.082
3
s3=0.2771
4
s4=0.0919
5
s5=0.0658
6
s6=0.1287
7
s7=0.1123
8
s8=0.0426
9
s8=0.0989
10
s10=0.1187
11
s11=0.166
12
s12=-0.0156
13
s13=0.2498
14
s14=-0.0119
15
s14=0.0792
Abundance
Mean Temperature

Figure S5:

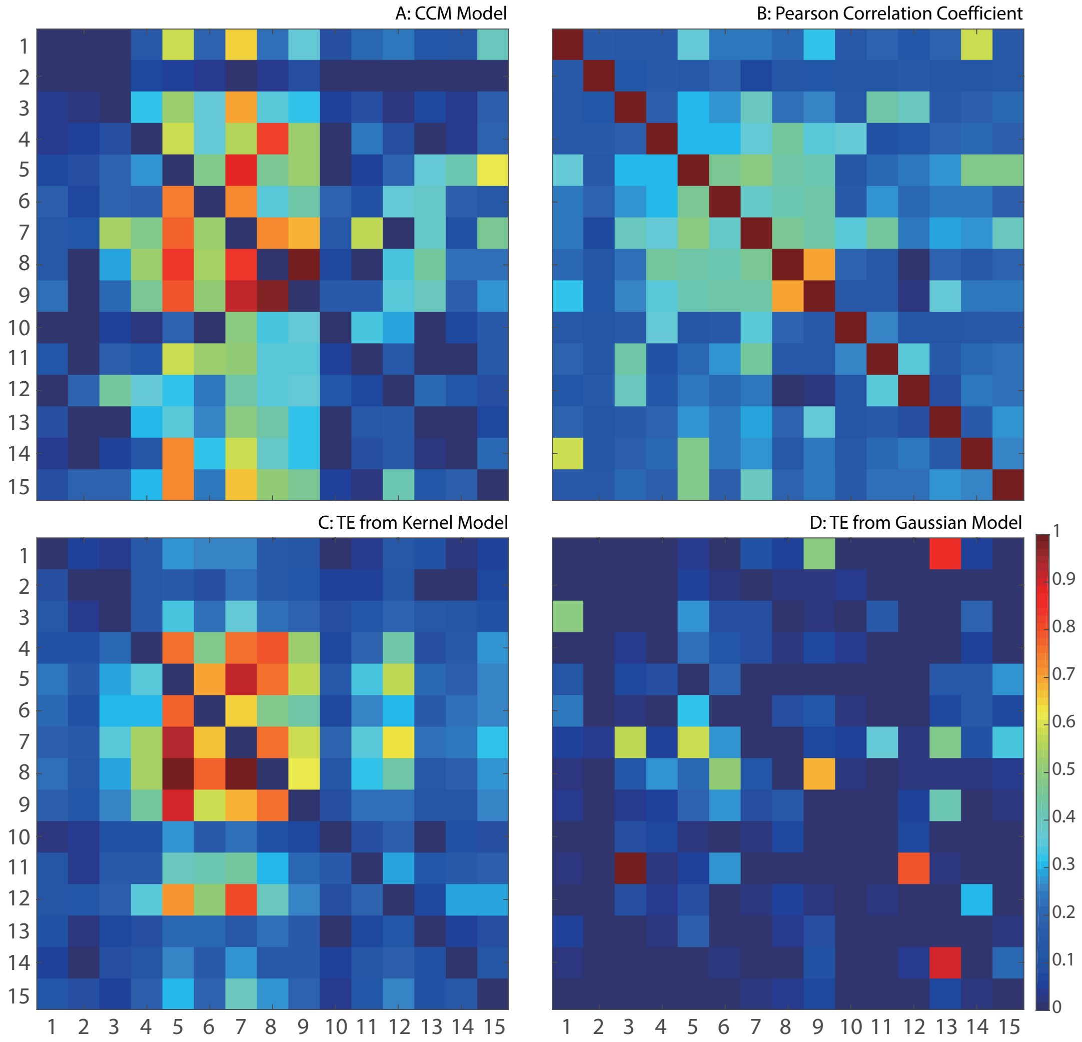
A: CCM Model
B: Pearson Correlation Coefficient
C: TE from Kernel Model
D: TE from Gaussian Model
1 2 3 4 5 6 7 8 9 10 11 12 13 14 15
0 0.1 0.2 0.3 0.4 0.5 0.6 0.7 0.8 0.9 1

Figure S6:

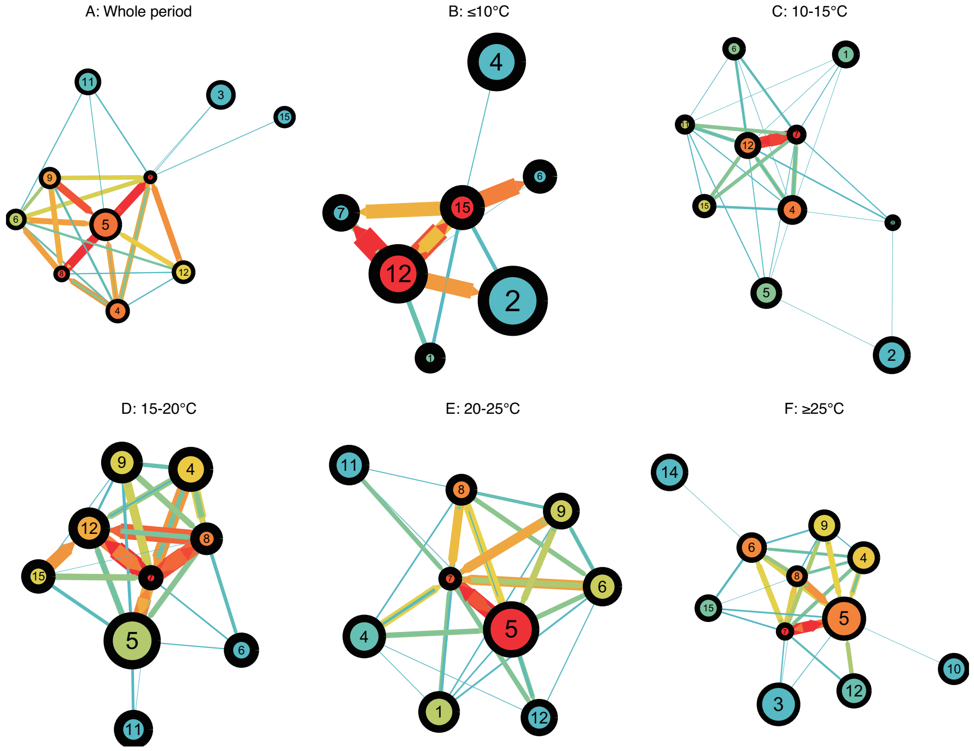

Figure S7:

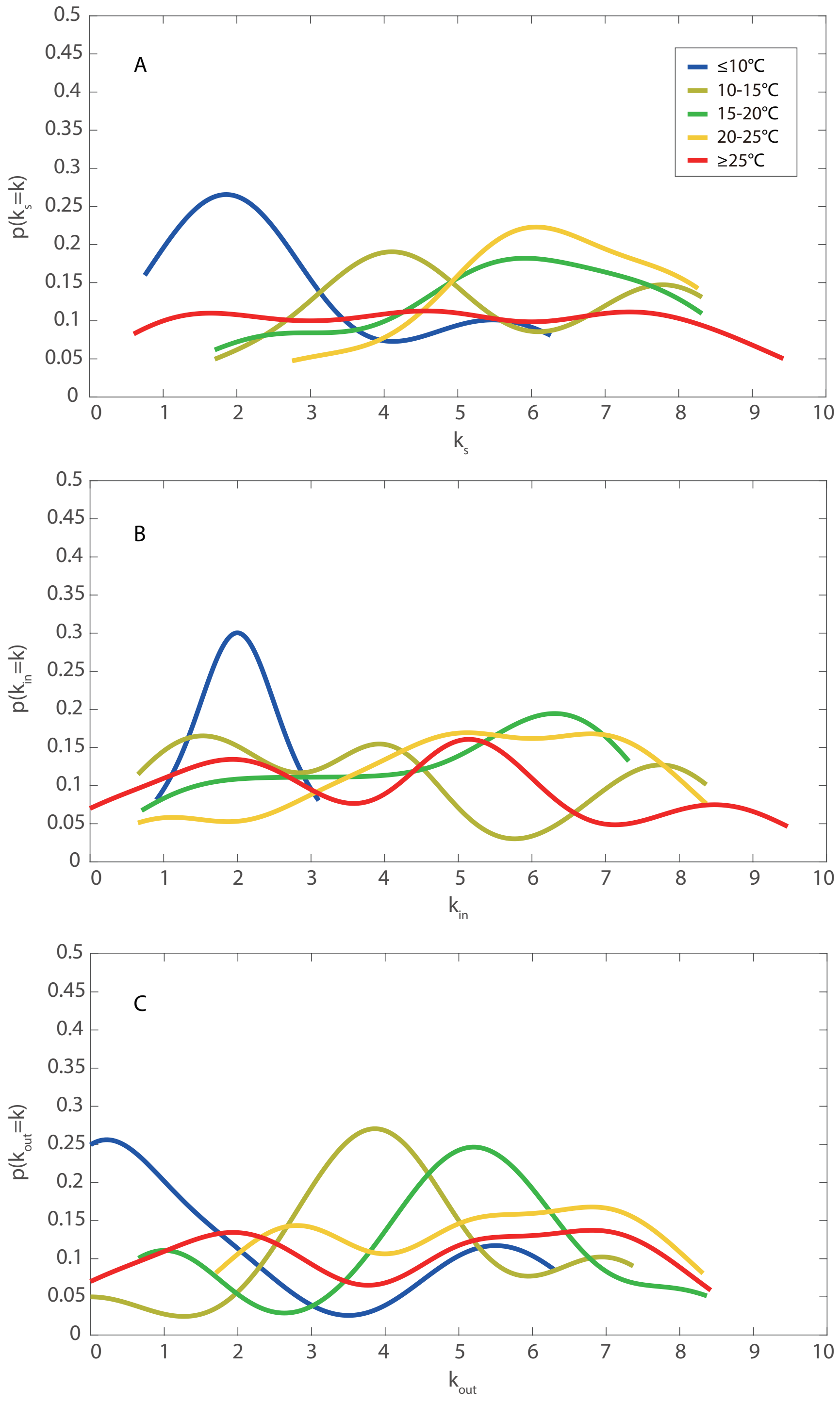

Figure S8:

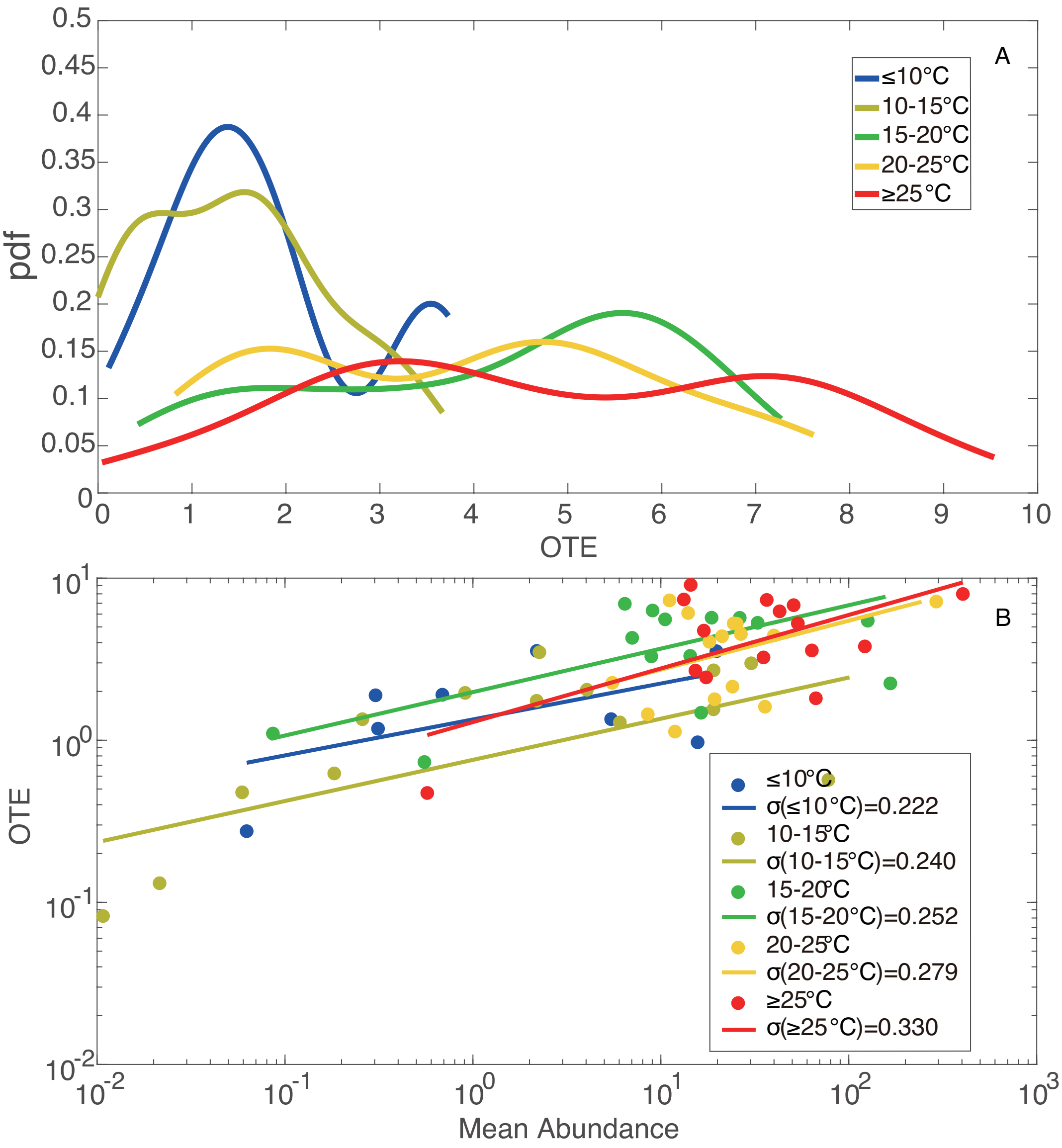

A
≤10°C
10-15°C
15-20°C
20-25°C
≥25°C
pdf
OTE
B
≤10°C
σ(≤10°C)=0.222
10-15°C
σ(10-15°C)=0.240
15-20°C
σ(15-20°C)=0.252
20-25°C
σ(20-25°C)=0.279
≥25°C
σ(≥25°C)=0.330
OTE
Mean Abundance

Figure S9:

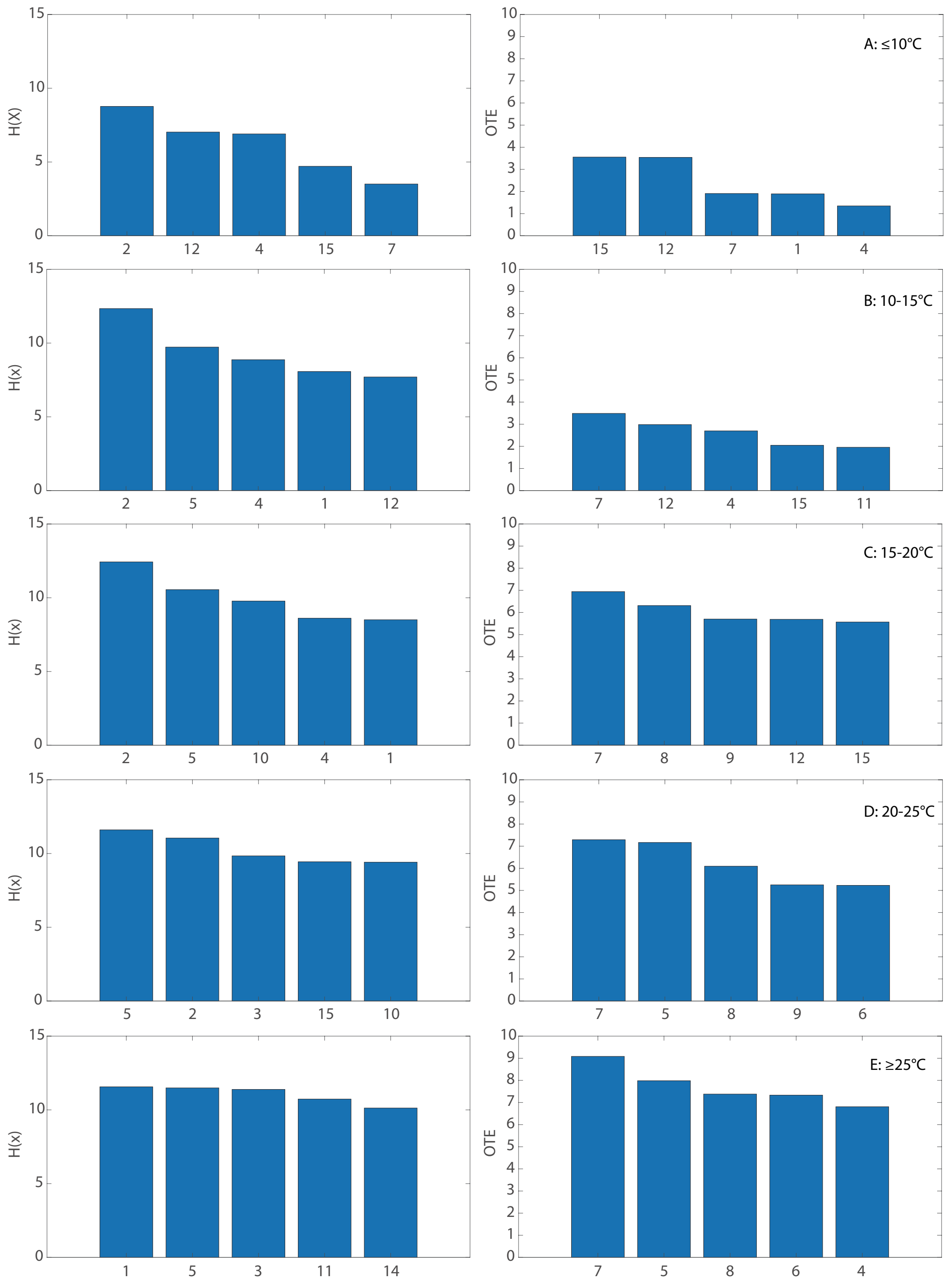

Figure S10:

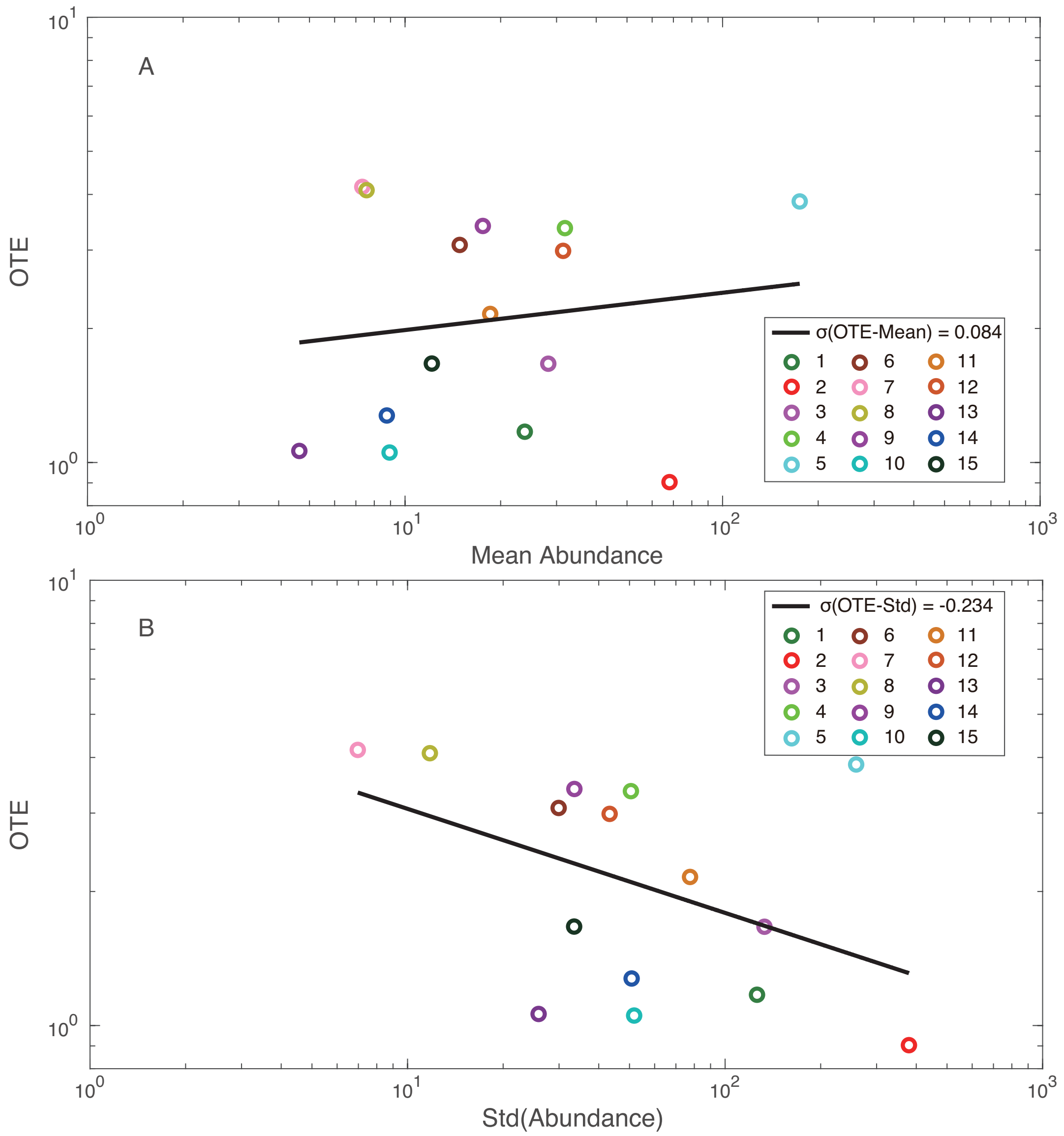
A
OTE
Mean Abundance
σ(OTE-Mean) = 0.084
B
OTE
Std(Abundance)
σ(OTE-Std) = -0.234

Figure S11:

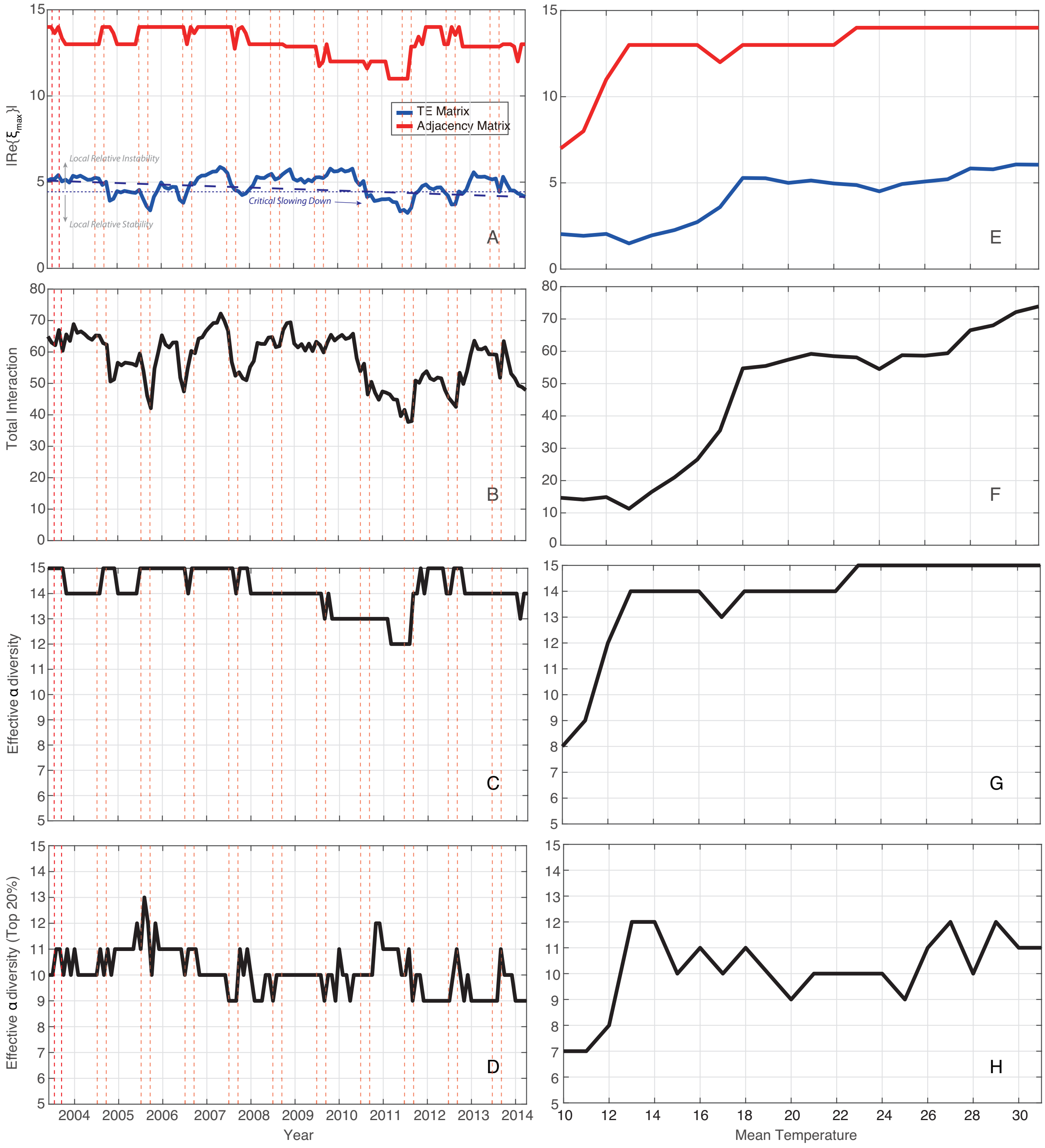

|Re{ξmax}|
TE Matrix
Adjacency Matrix
Local Relative Instability
Local Relative Stability
Critical Slowing Down
A
E
Total Interaction
B
F
Effective α diversity
C
G
Effective α diversity (Top 20%)
D
H
2004 2005 2006 2007 2008 2009 2010 2011 2012 2013 2014
Year
Mean Temperature

Figure S12:

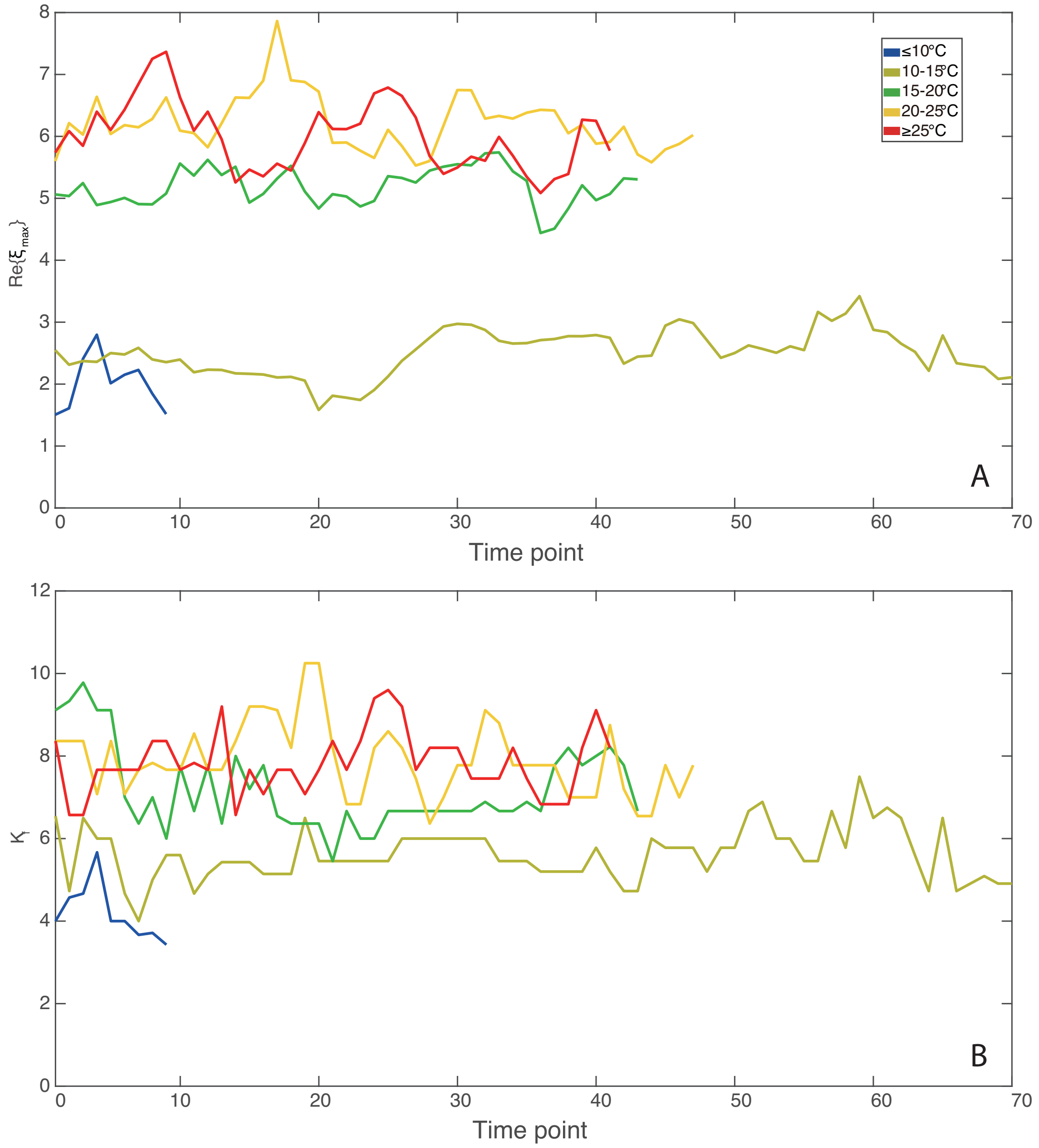

8
7
6
5
4
3
2
1
0
Re{ξ_max}
≤10°C
10-15°C
15-20°C
20-25°C
≥25°C
A
0
10
20
30
40
50
60
70
Time point
12
10
8
6
4
2
0
K_f
B
0
10
20
30
40
50
60
70
Time point

Figure S13:

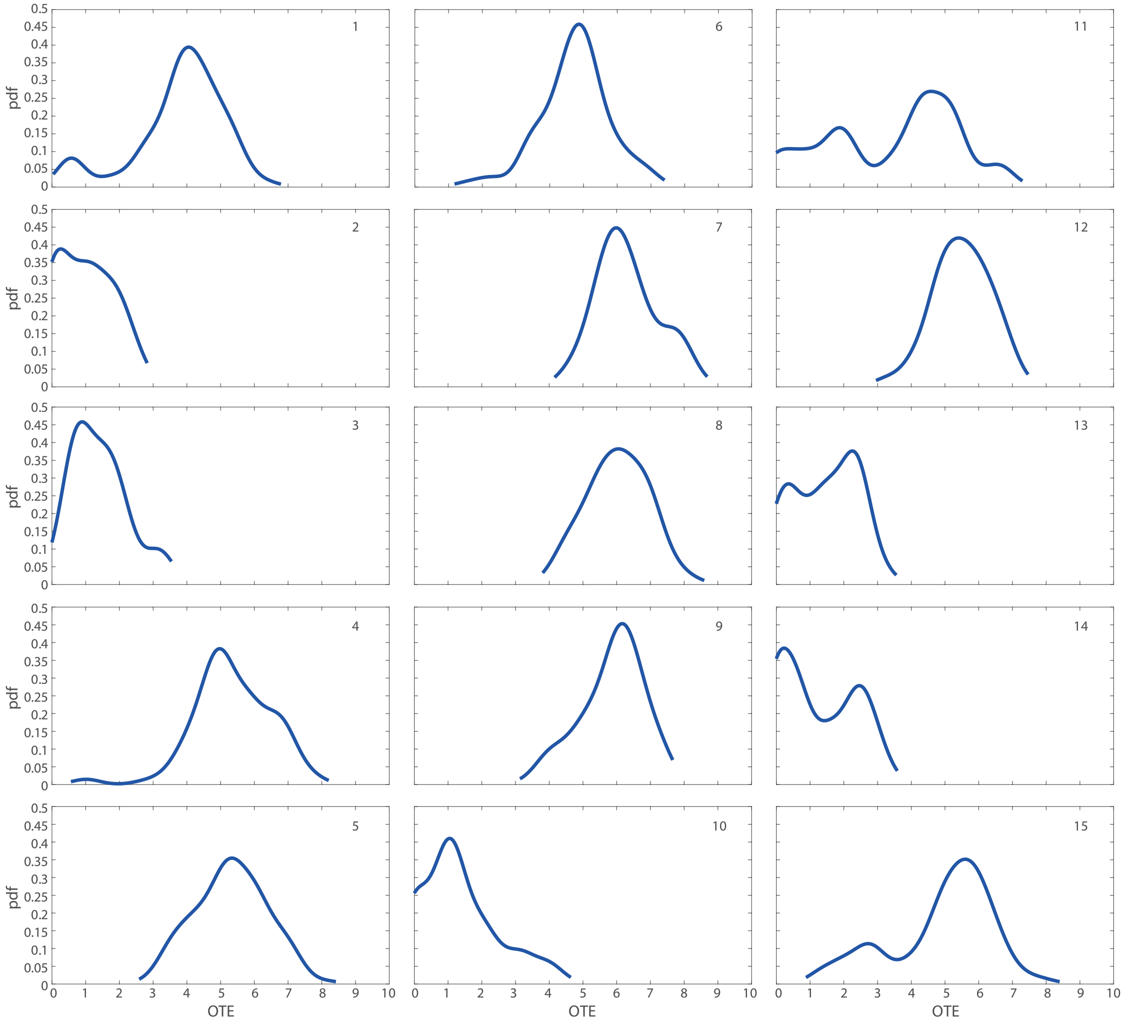

Figure S14:

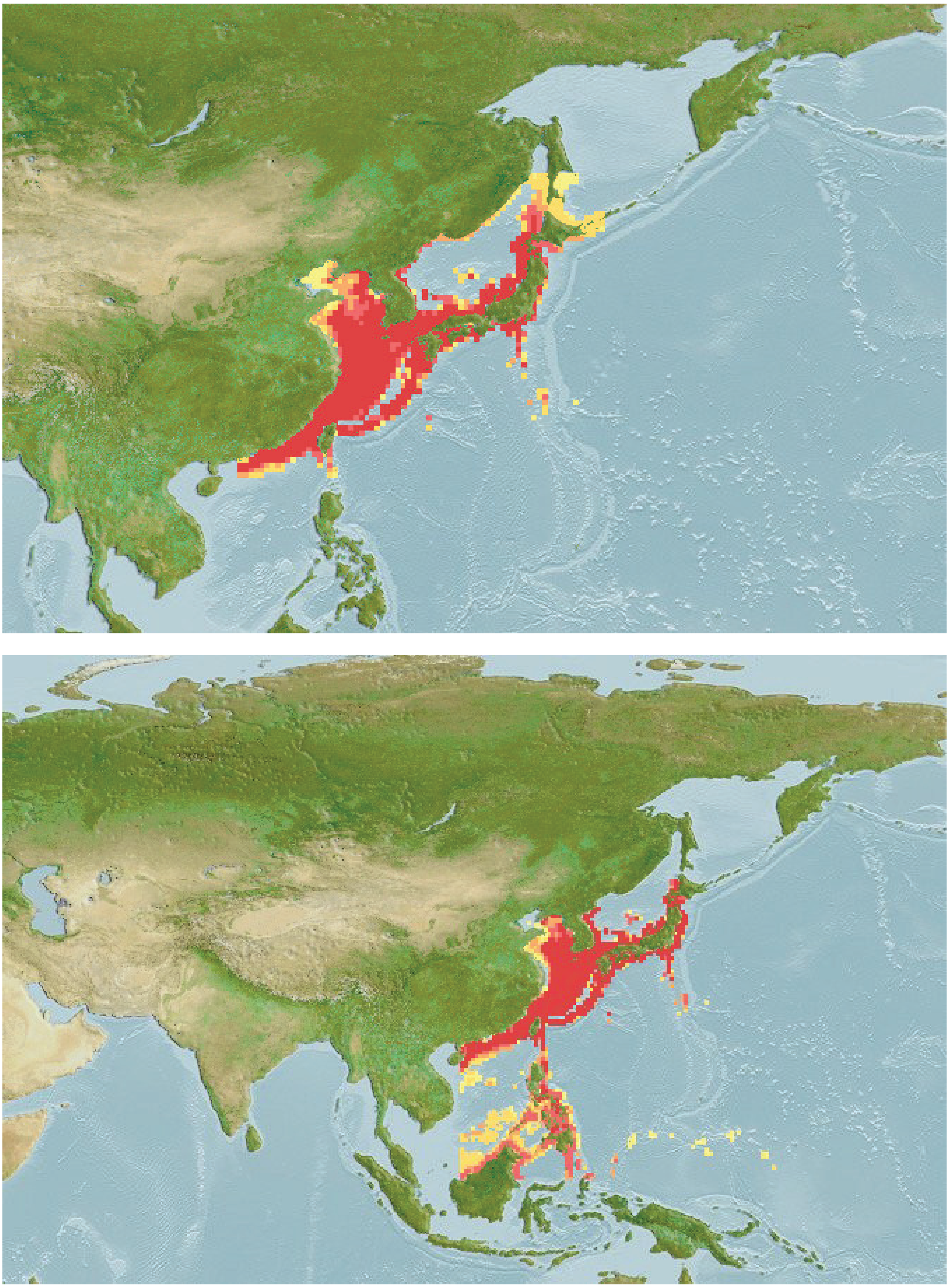

Figure S15:

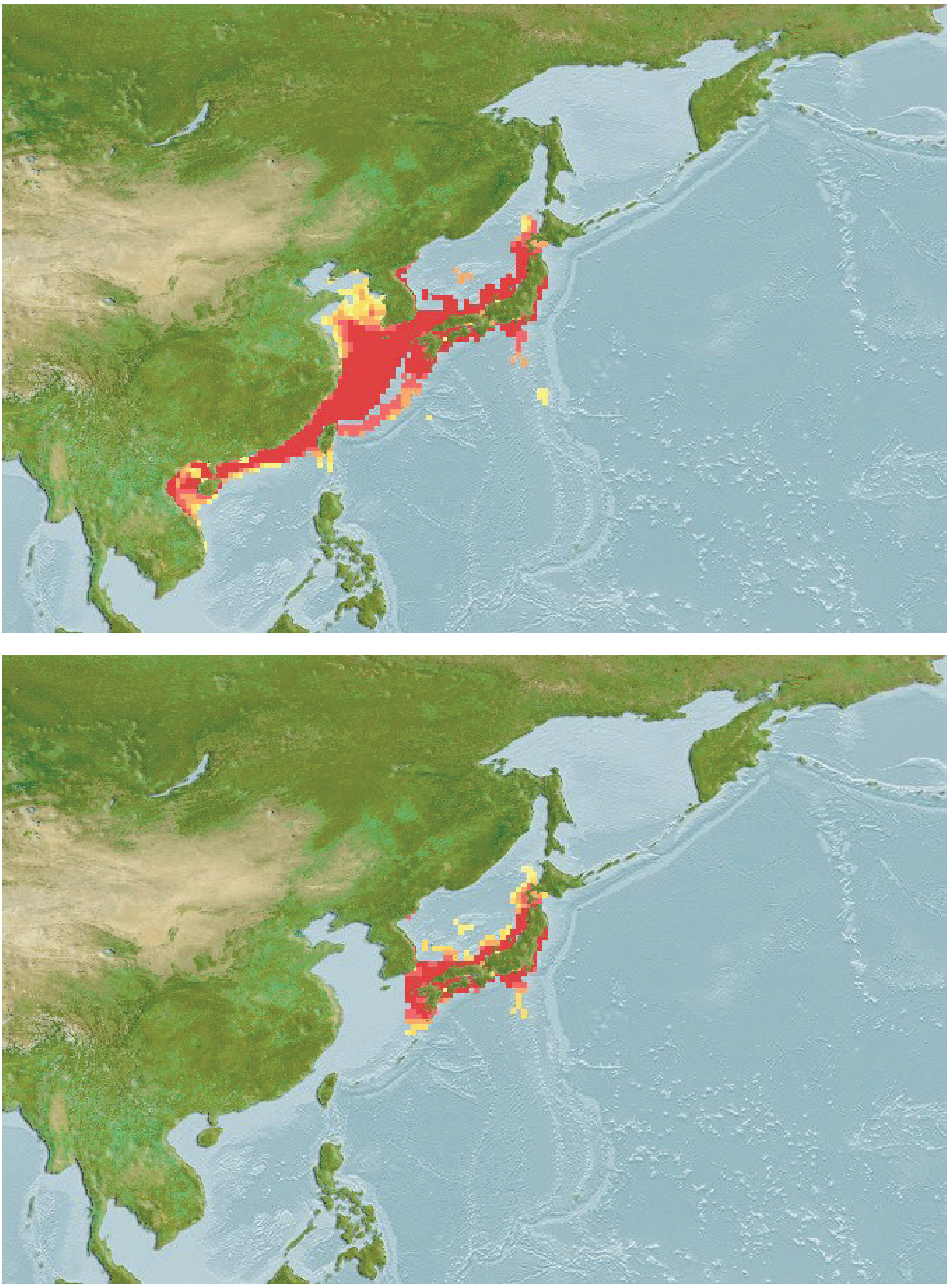

Figure S16:

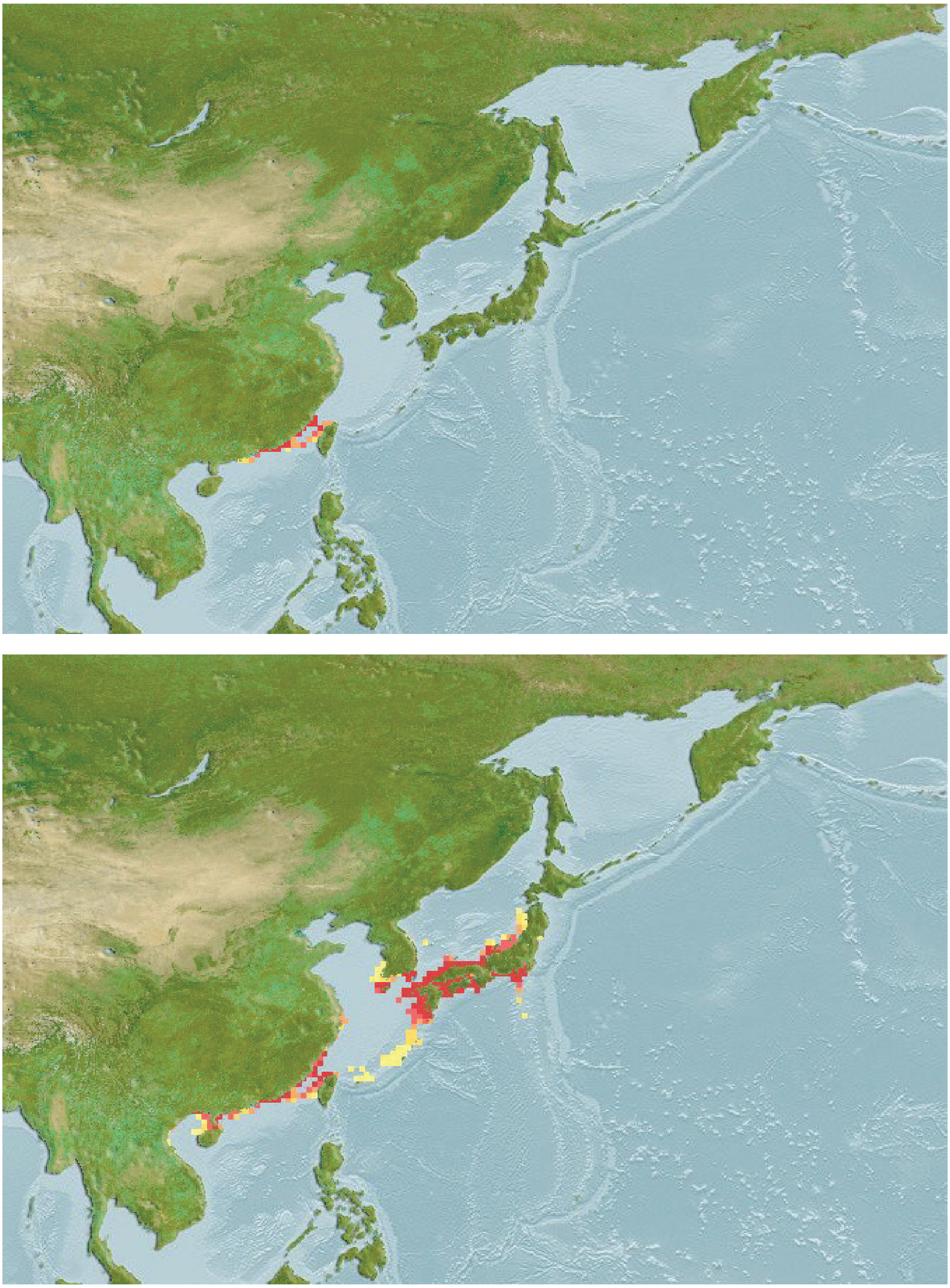

Figure S17:
